# Three annotated chromosome-level *de novo* genome assemblies of *Lomentospora prolificans* provide evidence for a chromosomal translocation event

**DOI:** 10.1101/2025.01.21.634120

**Authors:** Nina T Grossman, Yunfan Fan, Aleksey V. Zimin, Maggie P. Wear, Anne Jedlicka, Amanda Dziedzic, Livia Liporagi-Lopes, Winston Timp, Arturo Casadevall

**Author notes:** co-first author (Nina T Grossman, Yunfan Fan).

## Abstract

*Lomentospora prolificans* is a fungal pathogen responsible for serious, often fatal, illness in patients with compromised immune systems. Treatment is rarely successful because *L. prolificans* is inherently resistant to all major classes of antifungal drugs. In this study we publish three chromosome-level *de novo* genome assemblies, including the first complete-level assembly of *L. prolificans*, along with genome annotations. The L. prolificans genome is packaged in 11 nuclear chromosomes and one mitochondrial chromosome, has 36.7-37.1 Mb, and encodes for a putative 7357-7640 genes. The length and composition of contigs in one stain varied from those of the other two strains, supporting the hypothesis that a chromosomal translocation took place. These assemblies were confirmed with pulsed-field gel electrophoresis. The availability of more complete genomes will hopefully help the search for new antifungal drugs and provides insights into the evolutionary history of this pathogenic fungus.

## Introduction

*Lomentospora prolificans* is a filamentous fungal pathogen that causes disease primarily in severely immunocompromised patients, most often those with hematological malignancy and neutropenia (1). Though rare relative to other fungal diseases, these infections are very difficult to treat with current antifungal therapy, and, accordingly, disseminated infections have a mortality rate of 80% (2). The reason for this high frequency of antifungal treatment failure is that *L. prolificans* is intrinsically resistant to all three major families of systemic antifungal agents: azoles, echinocandins, and polyenes.

Thus far, relatively little research has been conducted on the causes for this remarkable drug resistance, and, of the three drug families, a mechanism of resistance has been proposed for only echinocandins. One major reason for this dearth of information is that, until very recently, no whole genome assembly was available for this organism. This changed in 2017 when Luo et al. published the first whole genome assembly of clinical *L. prolificans* isolate JHH-5317 (3). Sequenced with Illumina and Oxford Nanopore systems, this assembly consisted of 1625 contigs, of which 26 contained 98% of the total length of 37.63 Mb.

This whole genome sequence represents a massive resource for researchers wishing to study *L. prolificans*, and has already informed work on its mechanisms of resistance (4). However, in its current fragmented state and deriving from a single strain, the representativeness of the existing genomic data is unknown and this single genome sequence is an imperfect foundation upon which to build the entirety of *L. prolificans* molecular biology.

To generate a more complete, chromosome-level assembly of the *L. prolificans* genome, and of doing so with a greater diversity of strains, two additional *L. prolificans* isolates were sequenced and these sequences, as well as those generated in Luo et al., were each assembled *de novo*. To confirm the reliability of the resulting assemblies, pulsed-field gel electrophoresis was conducted. Accordingly, we have produced three *de novo* whole genome assemblies, one to complete level and two to near-complete chromosome level. These chromosome lengths have been verified by pulsed-field gel electrophoresis.

## Methods

### Strains

The strains used in this work were *L. prolificans* strains 3.1 from Christopher Thornton, ATCC strain 90853 and JHH-5317 from Sean Zhang. Conidia were obtained by inoculating frozen conidia into Sabouraud dextrose broth (BD, Franklin Lakes, NJ, USA), growing this at 30°C for 5-21 days shaking, then plating 2-3 ml onto potato dextrose agar. After 6-8 days of growth, these plates were covered in DPBS without magnesium or calcium and scraped. The resulting conidia in DPBS were strained through a 70 μm cell strainer, pelleted at 4000 rpm, washed three times, and stored at 4°C.

### DNA Isolation

DNA was obtained from *L. prolificans* strains by growing cultures in Sabouraud dextrose broth shaking at 30°C. Fungus was collected by centrifugation and washed with sterile water, then incubated in DNA isolation buffer (0.1 M Tris-HCl, 0.2 NaCl, 5 mM EDTA, 0.2% SDS) with 1 mg/ml proteinase K at 55°C overnight. Following incubation, fungus was subjected to bead beating on a vortex with 0.5 mm zirconia/silica beads for 10 min. Samples were then centrifuged at 13,000 RPM for 10 min at room temperature. Supernatant was removed, added to an equal volume of 1:1 phenol chloroform, vortexed and then centrifuged at 13,000 RPM for 5 minutes. The aqueous phase was transferred, added to an equal volume of chloroform, and vortexed. This was centrifuged at 13,000 RPM for 5 minutes, and the aqueous phase was removed. Twice the sample volume of molecular grade ethanol was added to the sample, which was then incubated at −20°C for at least 4 h. This was centrifuged at 13,000 RPM for 10 minutes at 4°C, following which the supernatant was discarded and 400 μl of 70% ethanol was added to the pellet. The sample was vortexed then centrifuged at 13,000 RPM for 10 min at 4°C. Supernatant was discarded, and pellet was washed with 400 μl of 100% ethanol, and vortexing and centrifugation were repeated. Supernatant was discarded, and pellet was allowed to air dry before being resuspended in molecular grade water and stored at −20°C.

### DNA sequencing

Oxford Nanopore Technologies (ONT) sequencing libraries were prepared from genomic DNA using the Ligation Sequencing Kit (SQK-LSK109) with the Native Barcoding Kit (EXP-NBD103) according to manufacturer specifications (Oxford Nanopore Technologies, Oxford, UK). Illumina sequencing libraries were prepared using the Nextera Flex DNA library prep kit (Illumina, San Diego, California) and sequenced on a MiSeq using v2 2×150 chemistry.

### Genome assembly and correction

ONT data were base-called and demultiplexed using Guppy v4.2.2 (Oxford Nanopore Technologies). Using reads greater than 3kb long, two assemblies were generated, one using Canu v2.1.1 and one using Flye v2.9, both on default settings with the estimated genome size set to 39 Mb (5–8). Nanopore reads were then aligned back to the assemblies using minimap2 v2.17. Alignments were used for correction with Racon v1.4.19 with settings -m 8 -x −6 -g −8 -w 500 (9,10). Further corrections were then made using medaka v1.2.1 with the r941_min_high_g360 error model (11). Illumina reads were trimmed using Trimmomatic v0.39 with settings LEADING:3 TRAILING:3 SLIDINGWINDOW:4:30 MINLEN:36 (12). Trimmed reads were then used to further correct the assemblies using Freebayes v1.3.4 in conjunction with the bowtie2 v2.4.2 aligner, both on default settings (13,14). Changes were made at loci where the alternative allele was both supported by more than five reads, and the alternative allele frequency was greater than 0.5. This Freebayes correction was performed iteratively until no changes could be made. To scaffold the contigs of the two corrected assemblies, Ragtag v1.0.2 was used on default settings (15). For 3.1 and 90853, the corrected assembly generated by Flye was used as the reference for Ragtag, while for 5317, the corrected assembly generated by Canu was used as the reference. The determination of the reference assembly was made based off of the telomere orientations of the resulting scaffolds.

Nanopore and trimmed Illumina reads were then aligned to the corrected assemblies using minimap2 and bowtie2 respectively, and contigs were broken at positions of zero coverage unless within 1kb of a contig end bearing telomere repeats (14,16). Mitochondrial contigs were identified with whole genome alignment to the *prolificans* reference genome from Luo et al. 2017 using Mummer v4.0.0 (3,17). One contig containing repeats of the entire mitochondrial genome was trimmed to a single mitochondrial genome, while other contigs comprised of incomplete mitochondrial sequence were discarded. Lastly, the mean coverage for each contig was calculated with nanopore and Illumina data separately. Contigs with coverage less than 90% of the median in both the nanopore and Illumina data were discarded.

### Analysis

Genome analysis was performed in Geneious (Biomatters, Auckland, New Zealand). Genomes were aligned to one another using LASTZ with default parameters (18,19).

Subtelomeres were discovered by aligning the final 10 kb of each contig terminating with repeats of the telomeric sequence TTAGGG with all others from its strain-specific assembly using Geneious aligner. The resulting alignment was then edited by hand. Additional subtelomeres were then located by aligning the resulting consensus to the full genome assembly using LASTZ.

### Pulsed Field Gel Electrophoresis

Conidia were inoculated into Sabouraud dextrose broth, grown with shaking at 30°C for ∼16 h, and used to generate protoplasts using minor modifications of published methodology (20). Briefly, fungal biomass was filtered through Miracloth (Millipore Sigma, Burlington, MA), washed with sterile water, and incubated at 30°C on a nutating mixer for between 4 h and overnight in OM buffer with 5% lysing enzymes from *Trichoderma harzianum* (Millipore Sigma). Contents were then split into sterile centrifuge tubes and overlaid with chilled ST buffer in a ratio of 1.2 ml fungal solution to 1 ml ST buffer. Tubes were then centrifuged at 5000 · g for 15 min at 4°C. Protoplasts were recovered at the interface of the two buffers and transferred to a sterile centrifuge tube, to which an equal volume of chilled STC buffer was added. Protoplasts were pelleted at 3000 g for 10 min at 4°C, following which supernatant was removed, and protoplasts were resuspended in 10 ml STC buffer. This was repeated two more times, with the final resuspension being performed with 200 ml GMB buffer. Plugs were then made and treated using methodology adapted from Brody and Carbon (21). Briefly, 200 ml of 1-2x 10^9^ protoplasts in GMB buffer were mixed with 200 ml 2 % low-melt agarose in 50 mM EDTA (pH 8) cooled to 42°C and pipetted into plug molds, which were then placed on ice for 10 min to solidify. Plugs were removed from the molds and incubated in NDS buffer with proteinase K at 50°C for 24 h, followed by three 30 min washes in 50 mM EDTA (pH 8) at 50°C, then stored in 50 mM EDTA (pH 8) at 4°C.

Plugs were inserted into gels made with SeaKem Gold agarose (Lonza, Basel, Switzerland), along with *Saccharomyces cerevisiae*, *S. pombe,* or *Hansenula wingei* size standards (Bio-Rad Laboratories, Hercules, CA) and clamped homogeneous electrical field (CHEF) electrophoresis was run in TAE on a CHEF-DR III (Bio-Rad Laboratories). Gels were run at an included angle of 106° at 14°C according to the following conditions: 0.8% agarose at 3 V/cm with a switch time of 500 sec for 48 h, at 14°C (figure 4a), 1% agarose at 2 V/cm with a switch time of 1800 sec for 100 h (figure 4b), and 0.8% agarose at 2 V/cm with a switch time of 1800 sec for 80 h (figure 4c).

After running, gels were stained with ethidium bromide and imaged using a ChemiDoc XRS+ gel imager (Bio-Rad Laboratories). Chromosome band lengths were estimated using Image Lab software (Bio-Rad Laboratories).

### RNA sequencing

Cultures were grown by inoculating conidia from strain JHH-5317 into 25 ml of RPMI to a concentration of 10^7^ conidia/ml, incubating the flasks at 37°C in 5% CO_2_ to facilitate germination for four hours, then shaking them for 13 h at 37°C in atmospheric conditions. Following this, 25 ml of RPMI containing 16 μg/ml of ITR, 16 μg/ml of VRC, 8 μg/ml of AMB or 2% DMSO by volume were added to each flask, for final concentrations of 8 μg/ml ITR, 8 μg/ml VRC, 4 μg/ml AMB or 1% DMSO. Flasks were shaken at 37°C for two hours, then strained through Whatman #2 filters (Cytiva, Marlborough, MA), washed twice with sterile Milli-Q water, added to 2 ml of Trizol Reagent (Thermo Fisher Scientific, Waltham, MA), flash-frozen and stored at −80°C. Three biological replicates were performed. All drugs were obtained from MilliporeSigma (Burlington, MA) and dissolved in DMSO.

Cell homogenization was performed in Trizol Reagent using silica spheres (Lysing Matrix C, MP Biomedicals, Irvine, CA) in a FastPrep 120 (MP Biomedicals), with 4 intervals at speed 6 for 30 sec. Homogenates remained on ice between shakes. Total RNA was purified using a PureLink RNA Mini Kit (Thermo Fisher Scientific), with the on-column PureLink DNase treatment, according to manufacturer’s instructions. RNA was quantified using a NanoDrop 1000. Quality assessment was performed by RNA ScreenTape Analysis in a TapeStation 2200 (Agilent Technologies, Santa Clara, CA).

Libraries for RNA-seq were prepared from 250 ng Total RNA using the Illumina TruSeq Stranded mRNA Library Prep kit, according to manufacturer’s Low Sample protocol (Illumina, San Diego, CA). Quality assessment of libraries was conducted by High Sensitivity ScreenTape analysis on a TapeStation 2200 (Agilent Technologies). Libraries were quantified by qPCR with the Kapa Library Quantification kit (Roche, Basel, Switzerland) in a StepOne Plus Real Time PCR System (Thermo Fisher Scientific). Libraries were diluted and pooled and a final quality assessment was performed using High Sensitivity DNA LabChip Analysis on a BioAnalyzer 2100 (Agilent Technologies). A paired end, 2 x 100bp, Illumina HiSeq 2500 run was performed at Johns Hopkins Genomics Genetic Resources Core Facility, RRID:SCR_018669.

### Genome annotation

RNA sequencing reads were aligned to each of the three genome assemblies using HISAT2 (22) and these alignments were input to Stringtie (23) to produce transcript assemblies. We then aligned a collection of 8390 Microascaceae family proteins available from Genbank, to each genome with NCBI tblastn, clustered the alignments and produced preliminary CDS features based on the local protein alignments with exonerate in protein2genome mode. We then filtered and reconciled the transcript and protein alignment features with gffcompare and gffread tools to produce final annotation in the GFF format. We then output the protein sequences and aligned them to the proteins in the UniProtKB database with blastp looking for a single best hit. Any protein that had a significant (e-value <10^-8) hit was annotated as being similar to that protein in UniProtKB, and its function was assigned in the GFF file. BUSCO was run on transcripts using gVolante (24,25).

## Results and Discussion

Two strains of *L. prolificans*, 3.1 and 90853, were sequenced using both Nanopore and Illumina for long and short reads. These reads, as well as the reads from the sequencing of strain JHH-5317 used in Luo et al., were then separately assembled *de novo*, resulting in genomes of 36.77, 37.05 and 36.73 Mb long, with 18, 13 and 12 contigs, for 5317, 3.1 and 90853, respectively (table 1). Each assembly contains one circular contig of ∼24 kb that aligned to the mitochondrial genome of the previously published *L. prolificans* genome.

**Table 1:**
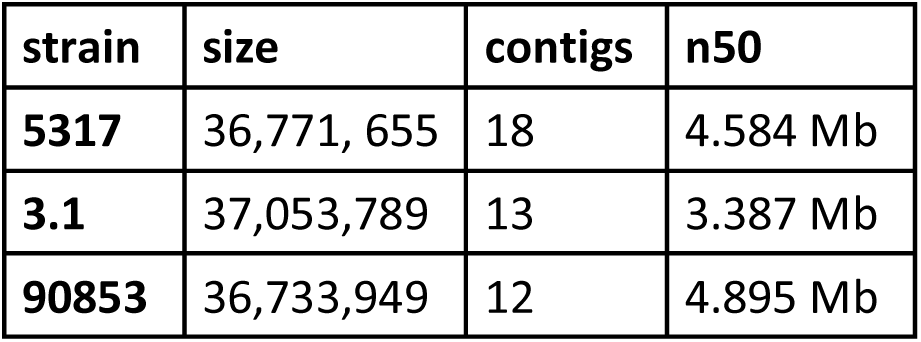
Length, contig number and n50 of the three *L. prolificans de novo* whole genome assemblies.

Of the 12 non-mitochondrial contigs in the 3.1 assembly, 10 have telomeric repeats (TTAGGG) on both ends, and the other two have telomeric repeats on one end (table 2). The contigs range in size from 6.89 to 1.22 Mb. In the assembly of 90853, nine of the 11 non-mitochondrial scaffolds have telomeric repeats on both ends, and another two have telomeric repeats on one end (table 3). The contigs range in size from 5.69 Mb to 1.22 Mb. Of the 17 non-mitochondrial contigs in the 5317 assembly, none have telomeric repeats on both ends, and only four have telomeric repeats on one end (table 4). These contigs range in size from 5.69 Mb to 7.27 Kb, with 12 contigs longer than 1 Mb.

**Table 2:**
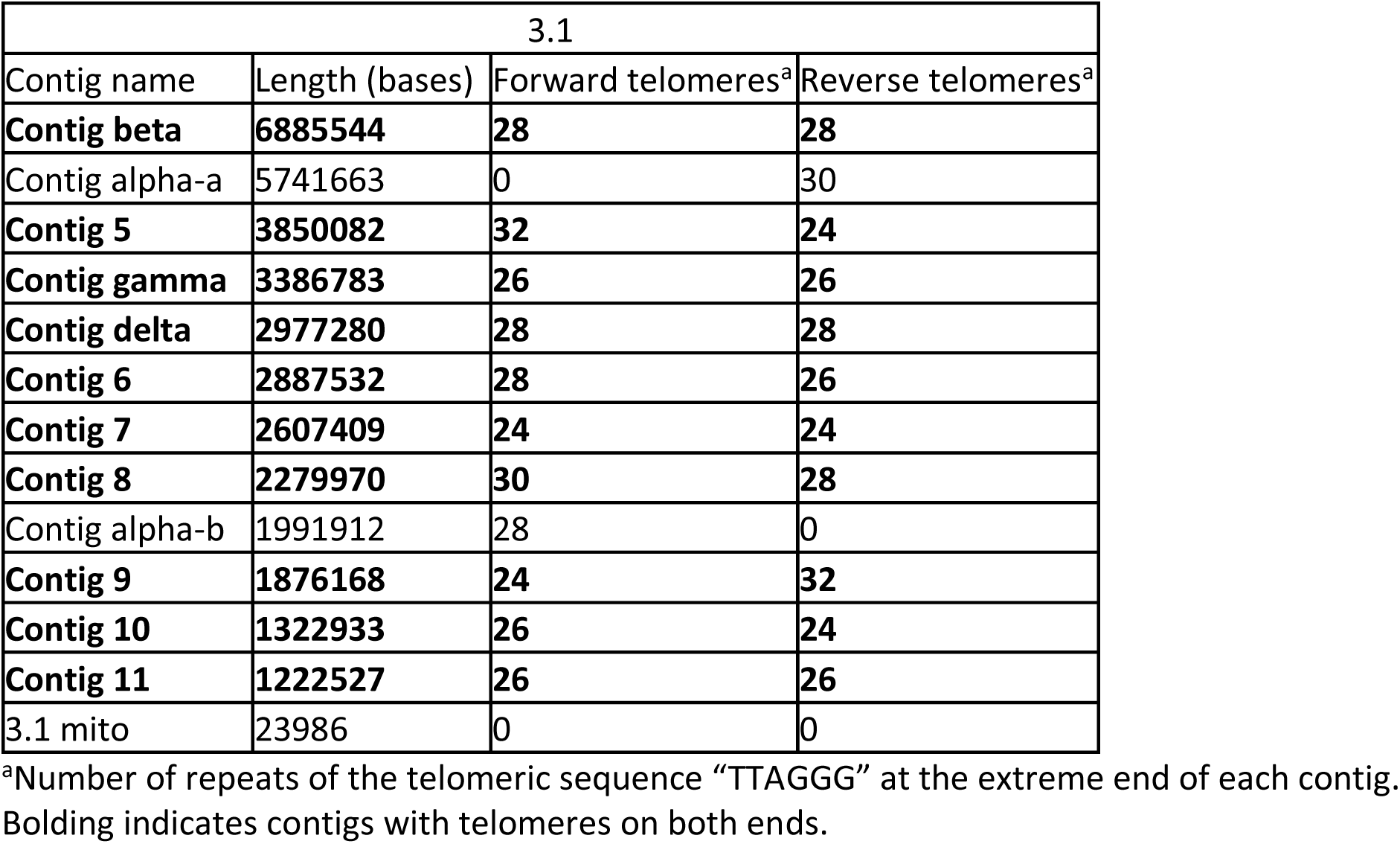
Contigs of the genome assembly of strain 3.1.

**Table 3:** Contigs of the genome assembly of strain 90853.

| 90853 |  |  |  |
| --- | --- | --- | --- |
| Contig name | Length (bases) | Forward telomeres<sup>a</sup> | Reverse telomeres<sup>a</sup> |
| Contig 1 | 5688415 | 30 | 0 |
| Contig 2 | 5146484 | 0 | 40 |
| **Contig 3** | **5112009** | **30** | **26** |
| **Contig 4** | **4894913** | **28** | **28** |
| **Contig 5** | **3779371** | **26** | **28** |
| **Contig 6** | **2935396** | **30** | **26** |
| **Contig 7** | **2533795** | **28** | **28** |
| **Contig 8** | **2283289** | **28** | **34** |
| **Contig 9** | **1781418** | **26** | **30** |
| **Contig 10** | **1336470** | **28** | **30** |
| **Contig 11** | **1218402** | **30** | **30** |
| 90853 mito | 23987 | 0 | 0 |

<sup>a</sup>Number of repeats of the telomeric sequence “TTAGGG” at the extreme end of each contig. Bolding indicates contigs with telomeres on both ends.

**Table 4:** Contigs of the genome assembly of strain JHH-5317.

| 5317 |  |  |  |  |  |
| --- | --- | --- | --- | --- | --- |
| Contig name | Length (bases) | Forward telomeres<sup>a</sup> | Reverse telomeres<sup>a</sup> | Forward subtelomere | Reverse subtelomere |
| *Contig 1* | *5,694,104* | *0* | *0* | *x* | *x* |
| *Contig 2* | *5,204,309* | *0* | *0* | *x* | *x* |
| *Contig 3* | *4,974,520* | *0* | *0* | *x* | *x* |
| Contig 4-a | 4,583,919 | 0 | 0 | x |  |
| *Contig 5* | *3,804,892* | *0* | *18* | *x* | *x* |
| *Contig 7* | *2,601,742* | *0* | *2* | *x* | *x* |
| *Contig 8* | *2,282,424* | *10* | *0* | *x* | *x* |
| Contig 6-a | 1,822,144 | 0 | 0 | x |  |
| *Contig 10* | *1,273,626* | *0* | *0* | *x* | *x* |
| *Contig 11* | *1,235,459* | *0* | *0* | *x* | *x* |
| Contig 9-a | 1,217,717 | 0 | 0 |  | x |
| Contig 6-b | 1,114,990 | 0 | 0 |  | x |
| Contig 9-b | 365,328 | 0 | 0 |  |  |
| Contig 4-b | 325,797 | 0 | 0 | x |  |
| Contig 9-c | 224,199 | 0 | 2 |  | x |
| 5317 mito | 24,023 | 0 | 0 |  |  |
| tig46 | 15,197 | 0 | 0 |  |  |
| contig 36 | 7,265 | 0 | 0 |  |  |

<sup>a</sup>Number of repeats of the telomeric sequence “TTAGGG” at the extreme end of each contig. Italics indicates contigs with subtelomeres on both ends.

### Inter-strain correspondence

To determine the correspondence of the genomes between the three strains, the assemblies were aligned to one another using LASTZ. As strain 90853 had the lowest number of contigs, these were used as the initial template and named contigs 1 through 11, starting with the longest contig. When the strain 5317 assembly was mapped to 90853, there was one 5317 contig mapped to one 90853 contig for 90853 contigs 1, 2, 3, 5, 7, 8, 10 and 11, and the matching 5317 contigs were named accordingly. For 90853 contigs 4, 6, and 9, two, two and three 5317 contigs, respectively, mapped to each 90853 contig, and these were named contig 4-a and 4-b, contig 6-a and 6-b and contig 9-a, 9-b, and 9-c (figure 1). Further evidence that contig 9-a and 9-b are in fact consecutive portions of the same chromosome is that, when mapped to 90853 contig 9, their meeting ends overlap approximately 11.6 kb with 96.8% pairwise identity. None of the other proposed consecutive contigs showed meaningful overlap.

**Figure 1:**
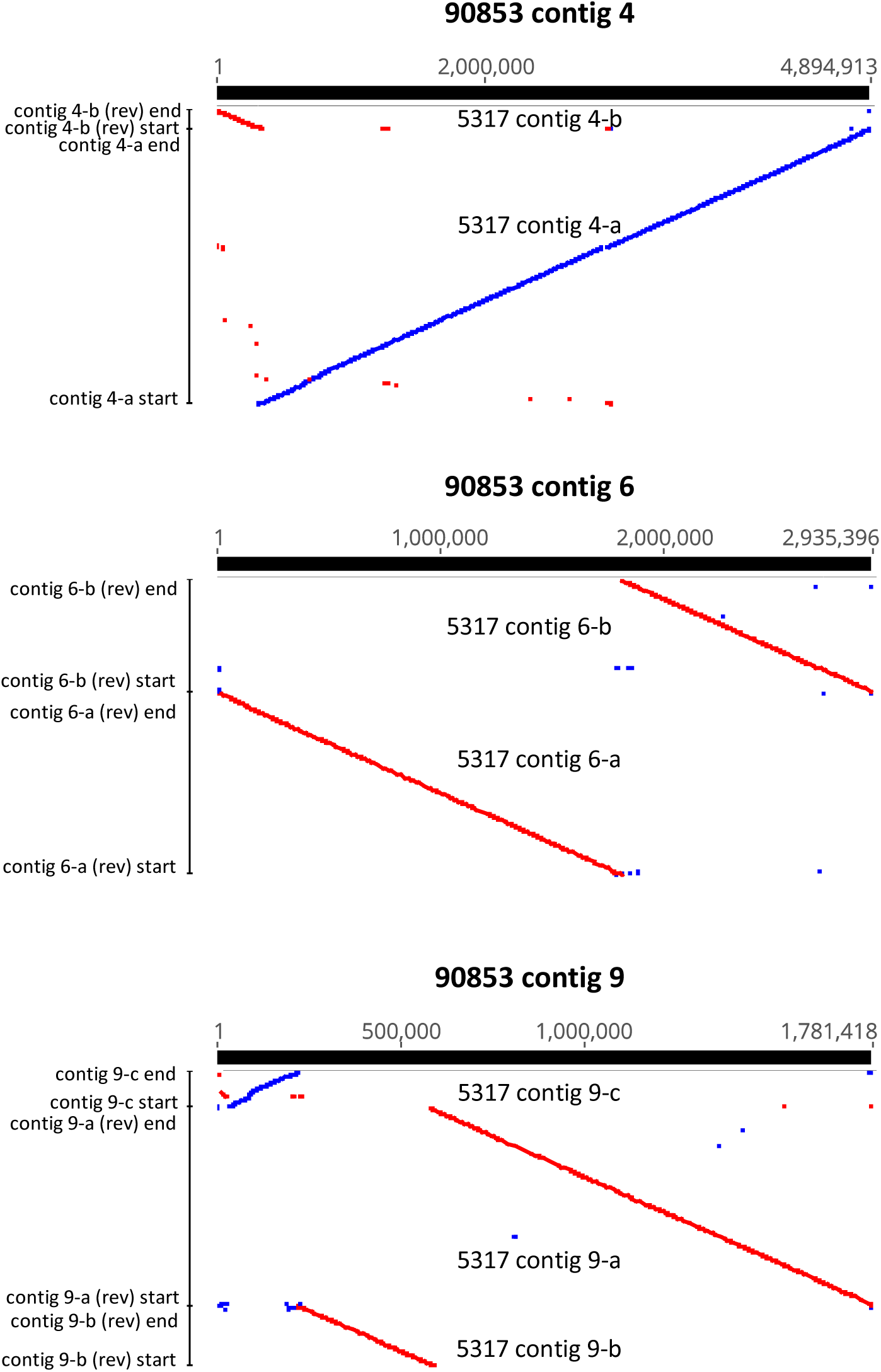
LASTZ alignments of 5317 contigs representing fragmented chromosomes against 90853 contigs representing complete chromosomes. Blue indicates forward sequence orientation and red indicates reverse sequence orientation.

The corresponding 90853 and 5317 contigs showed high pairwise identity, ranging from 98.1% to 99.1%, and high coverage, ranging from 92.0% to 99.5% for coverage of 90853 contigs by 5317 contigs and 96.1% to 99.4% for coverage of 5317 contigs by 90853 contigs (table S1). The strains’ mitochondrial genomes showed 99.4% pairwise identity and each mapped to the other for 99.1% coverage. The two smallest 5317 contigs did not align clearly to any 90853 contigs; only 52 of the 7,265 bases of contig 36 mapped anywhere on the 90853 genome, while tig46 mapped to multiple places on each of the 11 contigs of strain 90853 and likely represents repeated sequence.

When strain 3.1 was mapped to strain 90853, one 3.1 contig each aligned to 90853 contigs 5 through 11, showing between 98.1% and 99.1% pairwise identity, between 91.4% and 99.6% coverage of 90853 contigs by 3.1 contigs, and between 91.1% and 98.6% coverage of 3.1 contigs by 90853 contigs (table S1). The mitochondrial genomes showed 99.97% pairwise identity and 99.9% coverage for both strains. However, when the 3.1 assembly was mapped to 90853 contigs 1 through 4, evidence of a chromosomal translocation was observed. The first 5.4 Mb of 3.1 contig alpha-a mapped to the first 5.4 Mb of 90853 contig 1, but the remaining 0.32 Mb of 90853 contig 1 aligned to the last 0.32 Mb of 3.1 contig delta (figure 2). The first 2.6 Mb of 3.1 contig delta mapped to the first 2.6 Mb of 90853 contig 4, followed by the final 0.33 Mb of 3.1 contig alpha-a, and then the full 1.9 Mb of 3.1 contig alpha-b. This suggests an event in which 90853 contigs 1 and 4 each broke into two pieces and the four resulting pieces were incorrectly repaired, resulting in 3.1 contigs alpha-a and alpha-b, likely pieces of the same chromosome, and 3.1 contig delta.

**Figure 2:**
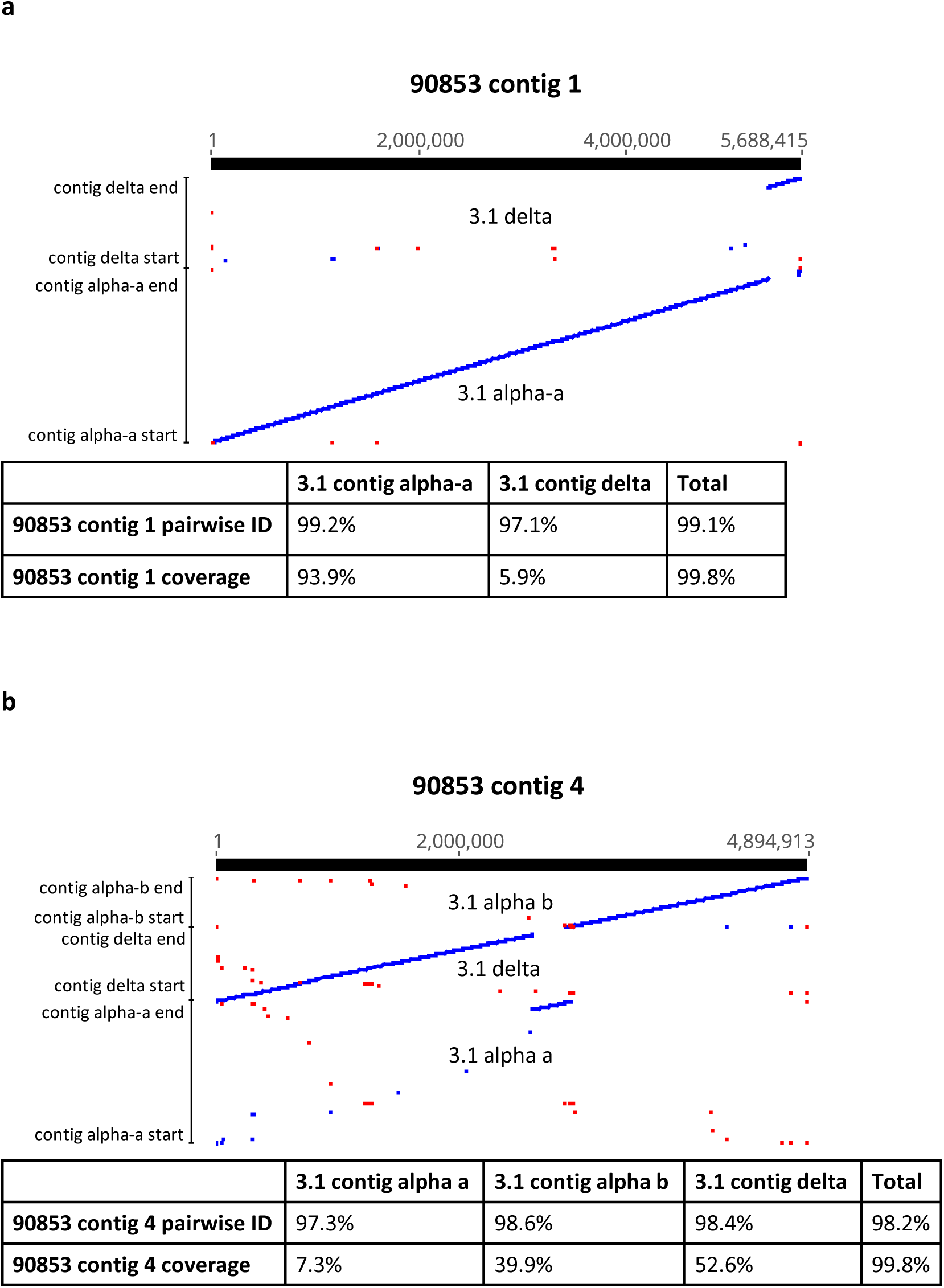
LASTZ alignments of strain 3.1 contigs alpha-a, alpha-b and delta to strain 90853 contigs 1 (a) and 4 (b). These contigs represent chromosomes that were involved in a chromosomal translocation. Pairwise identity and coverage are given below each alignment. Blue indicates forward sequence orientation and red indicates reverse sequence orientation.

These breakpoints are further supported by long read alignments, where 90853 nanopore reads aligned to contig 1 of the 90953 assembly showed no aberrant coverage near the breakpoint, while 3.1 nanopore reads aligned to the 90853 assembly showed coverage drops indicative of a rearrangement. Similarly, when nanopore reads of strain 3.1 are aligned to contig delta of the 3.1 assembly, no aberrant coverage patterns are observed, while alignment of 90853 nanopore reads to contig delta of the 3.1 assembly resulted in a coverage gap suggesting a structural variation exists at that locus. Taken together, the read alignment evidence argues against the possibility that these contig rearrangements are the result of any misassembly. The hypothesis that contigs alpha-a and alpha-b are two pieces of the same chromosome that were not joined by the assembler *in silico* is supported by the fact that the meeting ends of these two contigs share 15.1 kb of overlap with 96.8% pairwise identity. Additionally, each of the contigs has telomeric repeats only on its non-overlapping end, while all the other non-mitochondrial contigs of the strain 3.1 assembly have telomeric repeats on both ends, and are, accordingly, likely to be complete chromosomes.

A similar reconfiguration was seen with 90853 contigs 2 and 3 and 3.1 contigs beta and gamma, in which 3.1 contig beta seems to be made up of the first 4.2 Mb of 90853 contig 2 and the last 2.7 Mb of 90853 contig 3. 3.1 contig gamma, likewise, is made up of the first 2.5 Mb of 90853 contig 3 and the last 1.0 Mb of 90853 contig 2 (figure 3). These breakpoints are also supported by nanopore alignment evidence.

**Figure 3:**
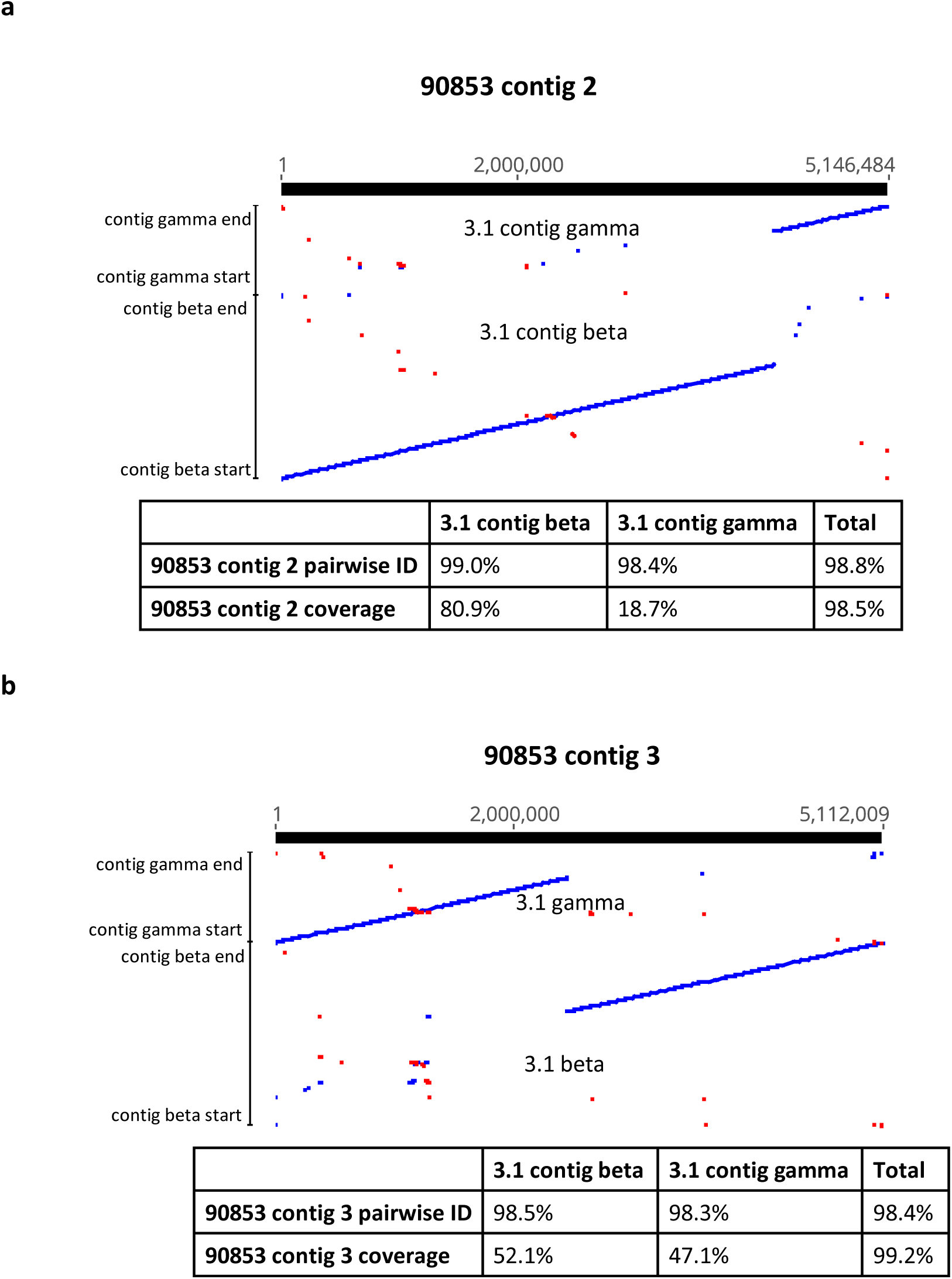
LASTZ alignments of strain 90853 contigs 2 (a) and 3 (b) and strain 3.1 contigs beta and gamma to one another. These contigs represent chromosomes that were involved in a chromosomal translocation. Pairwise identity and coverage are given below each alignment. Blue indicates forward sequence orientation and red indicates reverse sequence orientation.

### Subtelomeres

To determine whether the subtelomeric region of the *L. prolificans* genome has shared characteristics across chromosome and strains, the final 10 kb of contig ends containing telomeric repeats were aligned by strain. In strains 5317 and 3.1, all putative chromosome ends aligned with one another, resulting in consensus sequences 8,271 and 7,590 bases long with 95.4 and 95.8% pairwise identity, respectively (file S2 and S3).

No single sequence was conserved across the 20 putative chromosome ends of strain 90853, but three sequences were conserved across several putative chromosome ends each (file S4, file S5, file S6, and table S7). The shortest of these, consensus region 3, is, with some variation, the reverse complement sequence of a section of the longest, consensus region 1.

When each of the three strains’ conserved subtelomeric sequences were aligned to one another, those of 3.1 and 5317 showed 97.8% pairwise identity, while 90853 subtelomeric consensus region 2, shared by seven putative chromosome ends, mapped to both the 3.1 and 5317 sequences. The remaining subtelomeric sequences of 90853 did not map to those of the other two strains.

The presence of highly conserved subtelomeric sequences presented the opportunity to test subtelomeres as indicators of complete chromosomes on contigs where telomeres are not present. Accordingly, consensus sequences generated from each strain’s subtelomere alignment were mapped against that strain’s whole genome. In strain 3.1, this sequence mapped to each contig end that contained telomeres, and nowhere else in the genome. In strain 90853, subtelomeric consensus sequence 2 mapped exclusively to contig ends that contained telomeres, but only to eight of the total 20, which was in keeping with the lower level of subtelomeric conservation seen in this strain’s genome compared to the other two strains. The specificity of this conserved subtelomeric sequence to telomere-adjacent areas of the genome demonstrates that, in the absence of telomeres, searching for subtelomeric sequences in *L. prolificans* genomic assemblies is likely to be a specific, though not always a sensitive, means of identifying chromosome ends.

Mapping 90853 subtelomeric consensus region 1 back onto the assembly of strain 90853 produced less consistent results. Overall, it mapped to 14 regions in ten contigs that were within 20 kb of contig end and 12 regions more than 20 kb from an end. Accordingly, consensus with this sequence appears to be predictive of a subtelomeric region, but it is not unique to subtelomeres.

The principle of identifying chromosome ends using subtelomeric sequences was utilized for the strain 5317 assembly, which contains only four clusters of telomeric repeats. By mapping the conserved 5317 subtelomere sequence to the full assembly, 22 regions containing portions of the sequence were revealed, all located at the beginning or end of a contig. These were distributed such that the eight contigs that corresponded one-to-one with a full 90853 contig contigs contained subtelomeres on both ends, the six contigs that mapped to just one end of a full 90853 contig contained a subtelomere on just one end, and neither the two smallest contigs nor contig 9-b, which mapped to the middle of 90853 contig 9, had any subtelomeres. This provides compelling evidence for the correctness of genome assembly, given that subtelomeric sequences mapped only to the extreme ends of contigs. Additionally, it supports the identification of contigs or contig groups as chromosomes established through mapping the 5317 contigs to the 90853 assembly.

### Pulsed-field gel electrophoresis

To provide physical corroboration for the insights provided by the computational analysis of sequence data, pulsed-field gel electrophoresis was performed for the three sequenced strains, thereby visualizing their chromosomes. Examination of the resulting images shows nine bands for strains 90853 and 5317 ranging from 1.4 to 5.7 Mb in length, and 10 bands for strain 3.1 ranging from 1.2 to much greater than 5.7 Mb (above which estimation is not possible, as the longest ladder extends only up to 5.7 MB) in length (figure 4). The sum of the lengths of bands comes to 27.4, 27.5 and 28.2 Mb for strains 90853, 5317, and 3.1 (counting the large band of unknown size as 5.7 Mb), respectively. Given that these lengths are all approximately 9 Mb shorter than the lengths of the sequenced genomes, it is likely that at least one of these bands represents multiple chromosomes.

**Figure 4:**
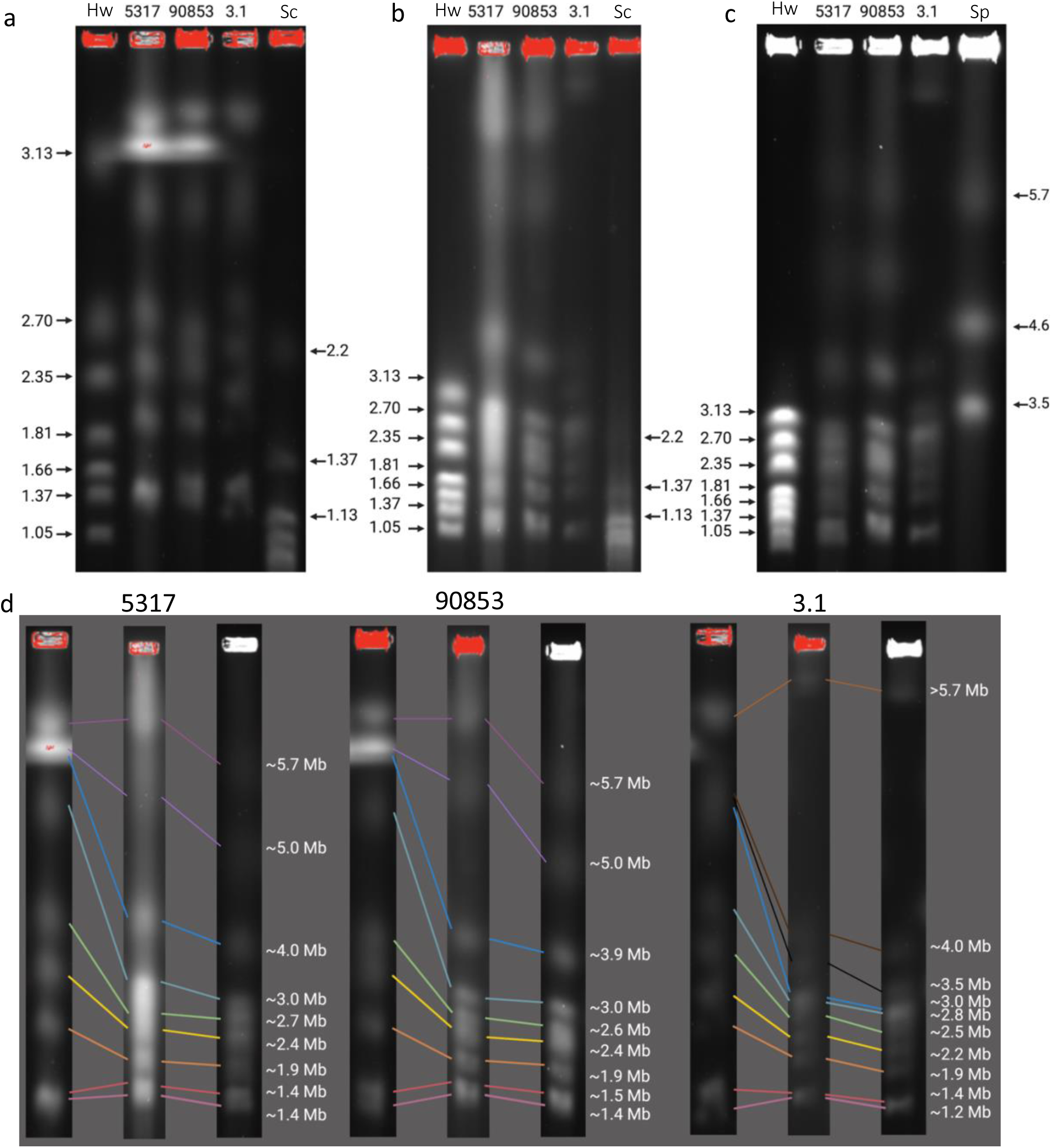
Pulsed field gel electrophoresis of *L. prolificans* strains. Chromosomal DNA from *H. wingei* (Hw), S. cerevisiae (Sc), and *S. pombe* (Sp) and *L. prolificans* strains JHH-5317, ATCC 90853 and 3.1, were run under three different conditions (a-c). Comparison of *L. prolificans* chromosomal bands with standard bands of known length allowed the estimation of the lengths of the *L. prolificans* chromosomes (d). See methods for condition details. Made using Biorender.com

It is notable that while strains 90853 and 5317 showed roughly the same banding pattern, strain 3.1 differed notably, with a band around 3.5 Mb where 90853 and 5317 show none, and no bands longer than 4.6 Mb until the longest band, high above 5.7 Mb, while 90853 and 5317 have three bands in this range (table 5). This corresponds to the chromosome sizes observed in the genome assemblies and confirms that the chromosomal translocation of 90853 contigs 1 through 4 into 3.1 contigs alpha through delta has actually occurred, resulting in chromosomes of distinctly different lengths.

**Table 5:** Correspondence of chromosome lengths in *L. prolificans* genome assemblies with chromosome lengths estimated by pulsed field gel electrophoresis.

|  | 90853 |  | 5317 |  | 3.1 |  |  |
| --- | --- | --- | --- | --- | --- | --- | --- |
| Chromosome | Length (Mb) | Matching band | Length (Mb) | Matching band | Length (Mb) | Matching band | Chromosome |
| 1 | 5.69 | ~5.7 | 5.69 | ~5.7 | 7.73 | >>5.7 | alpha |
| 2 | 5.15 | ~5 | 5.20 | ~5 | 6.89 | >>5.7 | beta |
| 3 | 5.11 | ~5 | 4.97 | ~5 | 3.85 | ~4 | 5 |
| 4 | 4.89 | ~5 | 4.91 | ~5 | 3.39 | ~3.5 | gamma |
| 5 | 3.78 | ~3.9 | 3.80 | ~4 | 2.98 | ~3 | delta |
| 6 | 2.94 | ~3 | 2.94 | ~3 | 2.89 | ~2.8 | 6 |
| 7 | 2.53 | ~2.6 | 2.60 | ~2.7 | 2.61 | ~2.5 | 7 |
| 8 | 2.28 | ~2.4 | 2.28 | ~2.4 | 2.28 | ~2.2 | 8 |
| 9 | 1.78 | ~1.9 | 1.81 | ~1.9 | 1.88 | ~1.9 | 9 |
| 10 | 1.34 | ~1.5 | 1.27 | ~1.4 | 1.32 | ~1.4 | 10 |
| 11 | 1.22 | ~1.4 | 1.24 | ~1.4 | 1.22 | ~1.2 | 11 |

### Genome completeness

To estimate the completeness of the three genome assemblies, BUSCO was used to search them for orthologs to the 3817 gene groups expected to be present as single-copy genes in *Sordariomycetes* (based on those found as such in ≥90% of the phylum) (25). The results were similar across the three assemblies, with 95.3-95.7% (3636–3653) of expected genes present as single copies, 0.5% (18–19) present in duplicate, 0.7-1.0% (26–37) present in a fragmented form, and 3.1-3.3% (120–125) missing entirely (table 6). This represents an improvement from the first published *L. prolificans* genome assembly that when compared to a set of 3725 expected gene groups, found 94.2% present either singly or in duplicate (3). Of the missing gene groups, 109 were shared across all three genomes. It is therefore likely that many of the “missing” gene groups are reflective of true absence from the *L. prolificans* genome, rather than failures in the sequencing or assembly.

**Table 6.**
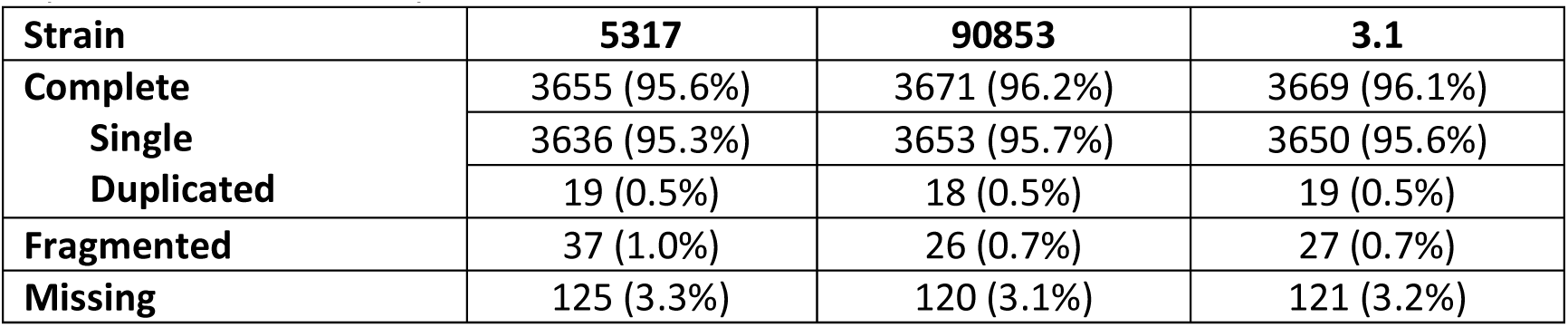
Results of BUSCO search of *L. prolificans* genome assembles for single-copy genes expected in *Sordariomycetes*.

### Annotation

The assemblies, in combination with ∼16.1 Gbp of sequence in Illumina RNA-seq reads from strain 5317 and a set of 8390 proteins from the family Microascaceae were used as evidence to generate gene annotation for the three *L. prolificans* strains (S8-10). The results are reported in table S11. These transcripts and the set of proteins were then used to produce annotations for the three strains, resulting in between 7559 and 7898 transcripts per strain and between 6927 and 7180 genes per strain (table S12). The BUSCO results of the transcripts are similar across the three strains, with 88.4-89.7% of expected genes appearing complete, ∼0.8-1.3% appearing in fragmented forms, and 9.2-10.3% missing (table S13).

### Conclusions

In this work, we determined that *L. prolificans* possesses 11 nuclear chromosomes made up of 36.7-37.1 Mb with 7357-7640 genes identified in each strain. The evidence of the telomeres, subtelomeres and pulsed-field gel electrophoresis provides compelling evidence that we have produced a complete-level assembly of *L. prolificans* strain ATCC 90853, and very nearly complete chromosome-level assemblies of strains JHH-5317 and 3.1. The BUSCO results show >95% of expected single-copy genes present in their complete forms, and of the 120-125 genes missing in the three strains, 109 are missing across all three, suggesting that a large portion of the missing genes are genuinely absent from the *L. prolificans* genome. Together, this supports the conclusion that these genome assemblies are complete.

The discovery of an apparent chromosomal translocation event in one of the three strains is intriguing and leads us to suggest that more *L. prolificans* strains be sequenced to provide a greater understanding of how anomalous such an event is in this species.

We believe the availability of complete, diverse and annotated chromosome-level genome assemblies of *L. prolificans* will be a valuable resource from which many new insights into *L. prolificans* biology can be gained.

## Data Availability

All sequence data are available in the Sequence Read Archive and Genbank, under BioProject PRJNA929059. All code used for analysis can be found at https://github.com/timplab/grossman_prolificans.

## Funding

YF was supported by the Canadian Institutes of Health Research Doctoral Foreign Study Award. A.C. was supported in part by NIH grants AI052733, AI15207, AI171093-01 and HL059842. NG was supported in part by 5-T32-AI-7417-22.

## Conflict of Interest

WT has two patents (8,748,091 and 8,394,584) licensed to ONT. WT and YF have received travel funds to speak at symposia organized by ONT.

---

**Table S1:** Similarity of three *L. prolificans* genome assemblies across orthologous contigs. NA where chromosomal translocation prevents one-to-one comparison. PI: pairwise identity, cov.: coverage, NA: not applicable.

|  | 90853 vs 3.1 |  |  |  | 90853 vs 5317 |  |  |  | 3.1 vs 5317 |  |  |  |
| --- | --- | --- | --- | --- | --- | --- | --- | --- | --- | --- | --- | --- |
|  | 90853 |  | 3.1 |  | 90853 |  | 5317 |  | 3.1 |  | 5317 |  |
|  | %PI | %cov. | %PI | %cov. | %PI | %cov. | %PI | %cov. | %PI | %cov. | %PI | %cov. |
| Contig 1 | NA | NA | NA | NA | 99.0 | 98.9 | 99.0 | 98.9 | NA | NA | NA | NA |
| Contig 2 | NA | NA | NA | NA | 99.0 | 99.5 | 99.0 | 99.4 | NA | NA | NA | NA |
| **Contig 3** | NA | NA | NA | NA | 98.8 | 96.9 | 98.7 | 98.9 | NA | NA | NA | NA |
| **Contig 4** | NA | NA | NA | NA | 98.6 | 97.7 | 98.7 | 97.5 | NA | NA | NA | NA |
| **Contig 5** | 99.1 | 98.7 | 99.1 | 97.2 | 99.1 | 98.5 | 99.1 | 97.9 | 99.1 | 98.6 | 99.1 | 99.4 |
| **Contig 6** | 98.8 | 96.6 | 98.8 | 98.0 | 98.6 | 98.7 | 98.7 | 98.4 | 98.7 | 98.6 | 98.7 | 97.1 |
| **Contig 7** | 98.8 | 99.6 | 98.9 | 97.2 | 98.8 | 98.9 | 98.8 | 96.5 | 98.9 | 99.2 | 98.9 | 99.2 |
| **Contig 8** | 98.6 | 97.6 | 98.8 | 98.0 | 98.8 | 98.0 | 98.8 | 98.1 | 99.3 | 99.2 | 99.3 | 99.1 |
| **Contig 9** | 98.6 | 96.3 | 98.5 | 91.1 | 98.4 | 97.4 | 98.4 | 97.0 | 98.8 | 93.2 | 98.9 | 97.7 |
| **Contig 10** | 98.1 | 91.4 | 98.0 | 92.9 | 98.1 | 92.0 | 98.0 | 96.1 | 98.9 | 95.3 | 98.9 | 98.6 |
| **Contig 11** | 98.9 | 99.0 | 98.9 | 98.6 | 98.8 | 99.2 | 98.8 | 98.1 | 98.8 | 98.5 | 98.8 | 98.5 |
| **mito** | 99.97 | 99.9 | 99.97 | 99.9 | 99.4 | 99.1 | 99.4 | 99.1 | 99.4 | 99.3 | 99.4 | 99.2 |

**Table S7:** Subtelomeric consensus regions from the genome assembly of strain 90853.

| **90853 consensus region** | **# chromosome ends** | **Length (bases)** | **pairwise identity** |
| --- | --- | --- | --- |
| **1** | 11 | 7,251 | 72.7% |
| **2** | 7 | 3,901 | 82.3% |
| **3** | 8 | 3,160 | 86.3% |

**Table S11.** Assembled transcript statistics for *L. prolifcans* genomes.

| **Strain** | **5317** | **90853** | **3.1** |
| --- | --- | --- | --- |
| **Number of transcripts** | 11640 | 12219 | 11930 |
| **N50 size of transcripts** | 4630 | 4185 | 4345 |
| **Total sequence in transcripts** | 37,058,901 | 36,362,687 | 36,441,817 |

**Table S12.**
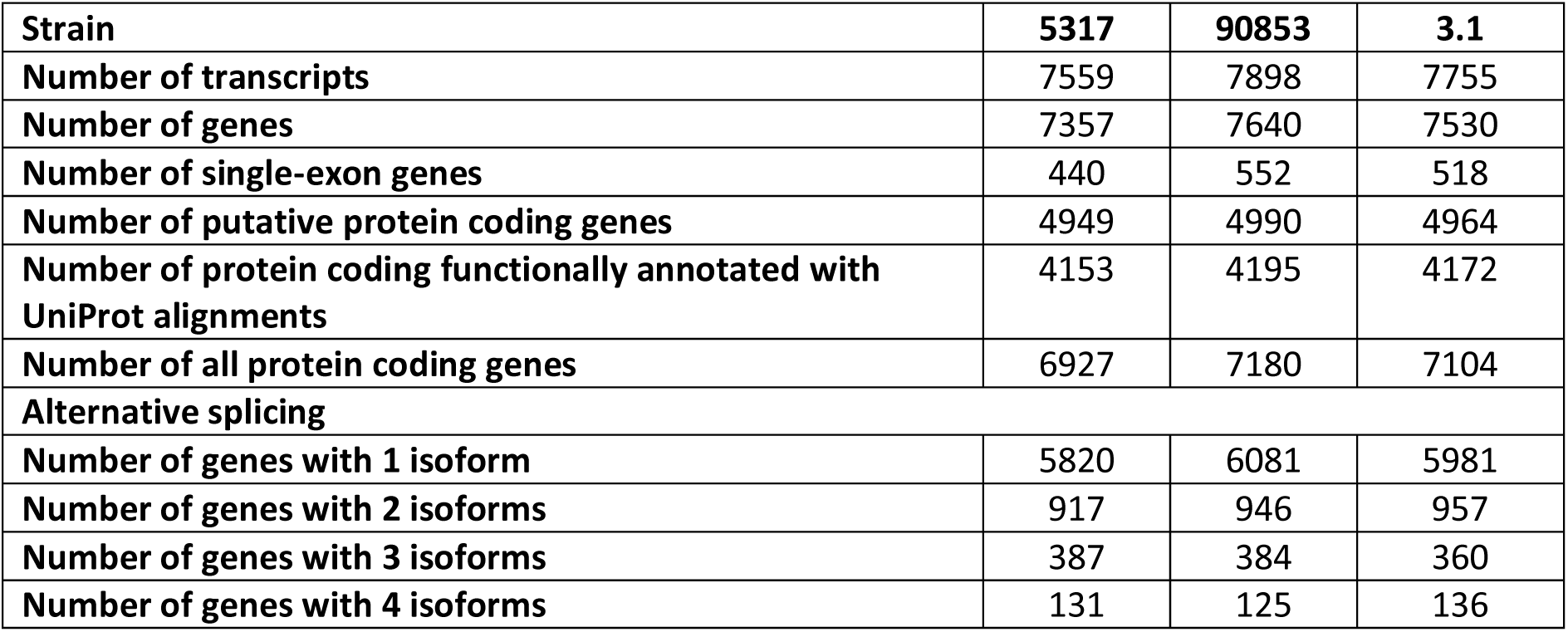
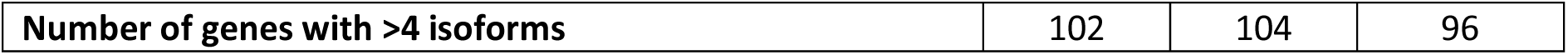
Annotation statistics for *L. prolifcans* genomes.

**Table S13.**
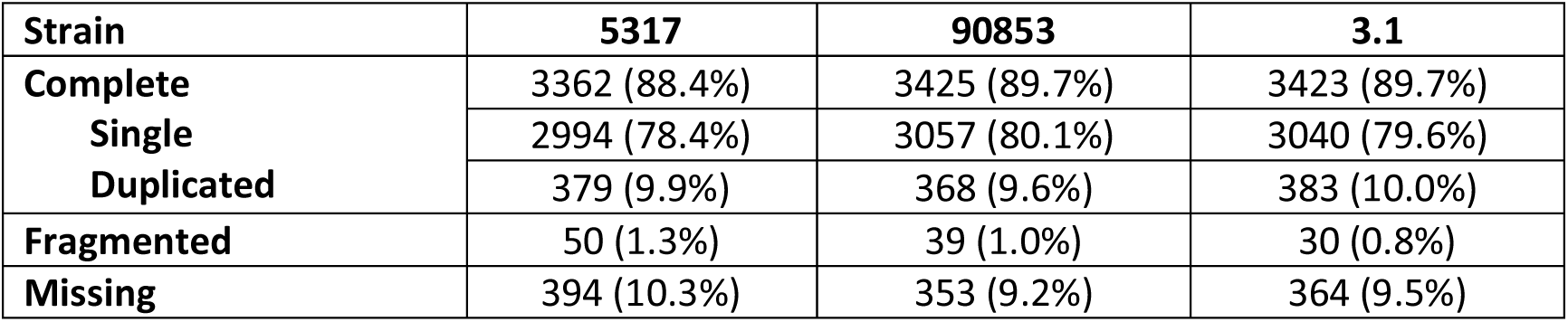
BUSCO results for *L. prolificans* transcripts.

